# Spatial overlap of sea ice-associated predators and prey in western Hudson Bay

**DOI:** 10.1101/2025.07.11.664287

**Authors:** C. Warret Rodrigues, A.E. Derocher, J.D. Roth, D. McGeachy, N.W. Pilfold

## Abstract

Spatio-temporal distribution of species shapes community structure and ecosystem function, yet the mechanisms driving biological hotspots remain unclear in dynamic environments like sea ice. We computed Getis-Ord (Gi*) hotspots based on four years of direct and indirect observations of polar bears (*Ursus maritimus*), Arctic foxes (*Vulpes lagopus*), ringed seals (*Pusa hispida*), and bearded seals (*Erignathus barbatus*) in western Hudson Bay, to identify spatial clustering and assess spatial relationships among these ice-associated species. We further mapped distribution hotspots of bear-hunting signs to examine predator-prey and intraguild relationships. Polar bears and bearded seals primarily used offshore areas, while Arctic foxes concentrated their activity on nearshore ice. Ringed seal lairs were distributed throughout the study area but mostly hauled out on nearshore landfast ice. The polar bear hotspot overlapped extensively ([30% - 49%]) with those of the three other species, and Arctic foxes had high overlap (50%) with ringed seals. In contrast, bearded seals and ringed seals had low overlap (18%), reflecting their different habitat preferences. Understanding current patterns in ice-associated species’ distributions and relationships is crucial to inform conservation actions and for predicting direct and indirect effects of Arctic warming. Our results help identify key ecological areas on sea ice and demonstrate how systematic collection of incidental observations can be combined to generate valuable ecological insights at low cost.

**Key-words:** Hotspot analysis, Predator-prey interactions, Commensalism, sea ice, *Ursus maritimus, Vulpes lagopus*, *Pusa hispida*, *Erignathus barbatus*

## Introduction

Spatial and temporal distribution of predators and their prey often drive interactions, affecting population dynamics of both predators and prey and thereby shape community structure and ecosystem functions [1–3]. While predators should preferentially forage in areas of higher prey availability [4,5], prey should avoid high-risk areas [6,7]. Prey can manage predation risk by minimizing the probability of encountering predators, or by coping with immediate threats to survival [8,9]. Depending on the outcome of the predators and prey interactions, their spatial distribution can correlate either positively (large spatial overlap) if the predator’s response prevails, or negatively (low spatial overlap) if the prey’s response prevails [10]. Relatively immobile prey, such as breeding animals after they establish their dens, lairs, or nests, will more likely respond reactively, rather than proactively [10,11], leading to large spatial overlap with their main predator.

Positive interactions between predators, such as commensalism or mutualism, have recently received recognition as a driving force of population dynamics and animal community structure [12]. In commensalist relationships, apex predators can benefit mesopredators by subsidizing them with carrion, with implications for mesopredator population dynamics, food-web structure and stability, and nutrient redistribution between ecosystems [13–16]. Polar bears (*Ursus maritimus*) are apex predators in the Arctic marine food web that depend on sea ice to travel, hunt, and reproduce [17,18].

Where sea ice is seasonal, like in Hudson Bay, polar bears undergo a period of hyperphagia in spring — peaking in May and matching ringed seal (*Pusa hispida*) pupping, mating, and molting periods — before ice break-up forces them to return to land where they fast until the ice reforms [19–21]. Polar bears primarily consume ringed seals throughout their range, but consume a variety of other prey, including bearded seals (*Erignathus barbatus*), harbor seals (*Phoca vitulina*), beluga whales (*Delphinapterus leucas*), and other marine mammals [18,21–23]. To maximize their energy intake, polar bears primarily consume the blubber of their prey, leaving the rest of the carcass [21,24] available for scavengers (Gamblin et al. 2025). For example, winter sea ice allows tundra predators and scavengers, such as snowy owls (*Bubo scandiacus*), ravens (*Corvus corax*), and Arctic foxes (*Vulpes lagopus*), to use marine resources to cope with terrestrial prey scarcity [25–27]. Because Arctic ecosystem productivity is low [28], carrion can be an important food source that supports the reproduction and winter survival of mesopredators [29–31]. However, the use of sea ice by terrestrial species and spatial relationships between ice-dependent and ice-facultative species are poorly understood.

The Arctic fox is the most well-known scavenger of polar bear kills [32]. In the Nearctic, Arctic foxes primarily feed on lemmings (Arvicolinae) year-round but they forage over sea ice when terrestrial prey are scarce [33–35]. In addition to scavenging on marine mammal carcasses, Arctic fox prey on newborn ringed seals [36,37], facing little interspecific competition in using marine resources, since their main competitor, the red fox (*Vulpes vulpes*), does not commonly venture onto the sea ice [38]. Arctic foxes start reproduction in spring and in years of low terrestrial prey availability, marine subsidies enhance reproductive success and increase adult survival, thereby stabilizing population sizes [25,30,39].

The abundant ringed seal and less abundant bearded seal are both broadly sympatric with a circumpolar distribution and peak abundance over continental shelf habitats with shallow waters and seasonal ice cover [40–42]. Ringed seals and bearded seals partition the niche space within the water column and through habitat preferences for haul out. Ringed seals primarily forage pelagically and semi-demersally on sea-ice associated prey, whereas bearded seals feed opportunistically on diverse benthic and pelagic prey [40,43]. Ringed seals maintain breathing holes throughout winter, allowing them to use landfast ice or pack ice [44–46]. Adults select stable consolidated ice with pressure ridges and other ice deformations on which snow accumulates and where they build subnivean lairs for protection against predators while resting, birthing, and nursing [20,45,47]. In contrast, bearded seals do not maintain breathing holes and pup on the surface of sea ice and thus rely on natural sea-ice openings that are prevalent in active pack ice [48–50].

The sea ice provides a platform linking all four species in the marine ecosystem.

It is a dynamic environment that undergoes major transformation throughout the year [51], but some features remain stable at coarse spatial scales, such as polynyas and flaw leads, which are formed by winds, currents, and tides. At a finer scale, however, these features vary temporally in shape and size and can modify the icescape on a scale of hours or days [52–54]. For species living in such shifting environments, the temporal variability in resources associated with a given geographical area will likely induce low persistence in their use of a particular area [55].

Aerial surveys have revealed general habitat preferences and variable densities of hauled-out ringed seals and bearded seals [56,57], but fine-scale distributional patterns remain unknown. Since most information comes from hauled-out individuals, little is known about the distribution of ringed seal birth lairs [46,58], with no information available from Hudson Bay. Polar bear space use has been investigated in relation to prey in several parts of its range [46,59,60], but its spatial overlap with scavengers remains largely unexplored. The use of sea ice for foraging and long-range movement by Arctic foxes is well documented [33,34,39,61], yet detailed sea-ice habitat use remains unknown. To date, no study has examined the spatial relationships among polar bears, Arctic foxes, and their pinniped prey.

Our objective was to identify distribution hotspots and spatial relationships among polar bears, Arctic foxes, ringed seals, and bearded seals in the sea ice environment. We hypothesized that both polar bears and Arctic foxes, which face limited interspecific competition, maximize their spatial overlap with resources. We further hypothesize that persistence in space use is low between years, due to the dynamic nature of sea ice. We predicted high spatial overlaps between polar bears and all three other species, a large overlap between Arctic foxes and ringed seals, and a low overlap between Arctic foxes and bearded seals, and between ringed seals and bearded seals reflecting their distinct habitat preferences. Finally, we did not expect high overlap between the yearly hotspots of polar bears and ringed seals based on the temporal dynamics of sea ice.

## Methods

### Study area

The study area encompassed 16,704 km^2^ of sea ice on western Hudson Bay, primarily to the north and east of Churchill, Manitoba, Canada, extending from ∼58.3°N to 59.5°N and 94.3°W to 92.5°W (Fig. 1). Hudson Bay is a shallow inland sea characterized by counterclockwise currents and annual ice [62]. The ice covers >90% of the Hudson Bay’s surface from December to May, when it starts breaking up. Sea ice starts to decay in the south and east under the influence of the prevailing northwestern winds, until the Bay becomes ice-free in July [51].

**Fig. 1.**
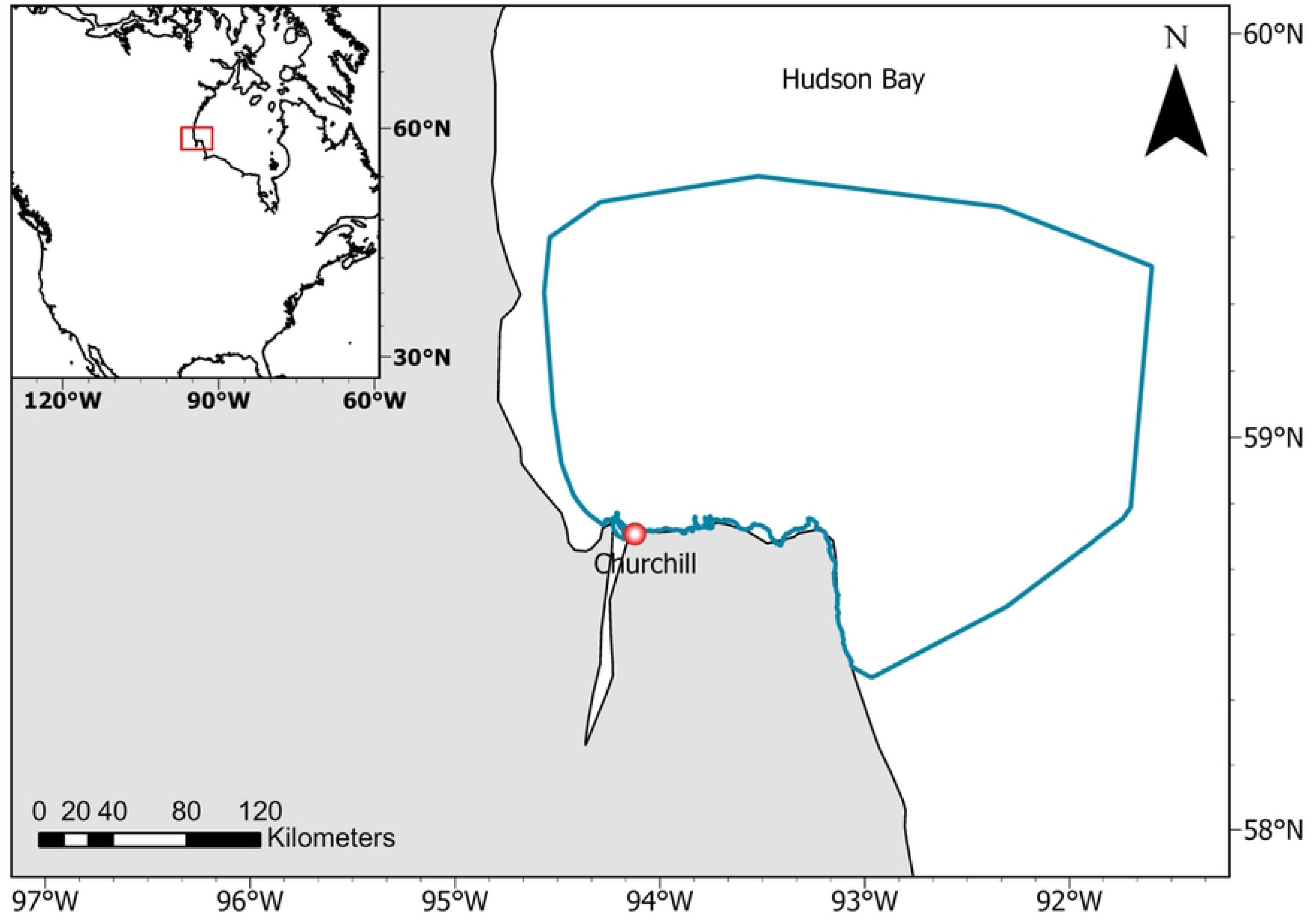
Study area (blue bounding box) delimited by minimum convex polygon on all helicopter locations pooled over 2019 to 2024.

### Flight survey

Using a Eurocopter AS350 B2 helicopter, we surveyed large areas of sea ice focusing on polar bear captures while tracking individual bears to study their population ecology. During the search, we systematically recorded the coordinates and characteristics of direct observations and signs (tracks) of polar bears, Arctic foxes, and common ice-associated species, with CyberTracker (cybertracker.org, Noordhoek, Cape Town, South Africa) on a Samsung A2 tablet (Samsung, South Korea). The surveys were interrupted in 2020 and 2021 due to pandemic restrictions, which resulted in a dataset spanning 4 years (2019, 2022-2024). Flights occurred when weather conditions had minimal cloud cover and winds < 30 km/h. Helicopter location was recorded every 5 seconds in 2019 and every second in the other years. Typical flight altitude was 75-150 m, and survey duration was 11.25 ± 2.75 days/year [8-14 days] starting on a mean of April 21 [April 16-25] and ending on a mean of May 1 [April 29 – May 5] (Table 1, S1 Table). The number of observers ranged from two to four in addition to the pilot.

**Table 1.** Sampling effort each year during helicopter surveys for signs of polar bears, seals, and Arctic foxes on the sea ice of western Hudson Bay.

| Year | Start date | End date | Number of survey days | distance flown (km) | Time flown (min) |
| --- | --- | --- | --- | --- | --- |
| 2019 | 20-Apr | 30-Apr | 9 | 3753 | 7144 |
| 2022 | 21-Apr | 05-May | 10 | 5336 | 6726 |
| 2023 | 25-Apr | 03-May | 9 | 4858 | 6474 |
| 2024 | 16-Apr | 29-Apr | 11 | 5322 | 9569 |

Observations recorded in 2019 included individual polar bears, hauled-out seals (species and number of individuals), hunting attempts (i.e., excavation of ringed seal subnivean lairs or breathing hole identified through bear track presence from the air; Pilfold et al. 2014), and successful predation events of polar bears on seal (hereafter “seal kills”) identified by the presence of carcass and blood [63]. The carcasses of seals could not always be identified to species level; therefore, all kills were pooled. In 2022, we added polar bear and Arctic fox tracks.

### Data standardization

We first thinned helicopter tracks to one location per minute to standardize across years. The survey area was defined by a 3.125 × 3.125 km grid (to match the sea ice data) within a bounding box, which we delineated using a minimum convex polygon encompassing all helicopter tracks across years. Because survey flights were regularized line transects, we accounted for uneven spatial survey effort by normalizing species observations per grid cell by dividing the number of observations by the number of helicopter locations + 1 in that cell.

### Hotspot analysis

We conducted hotspot analyses in R v. 4.2 (R Core Team 2024) using RStudio v.2024.12.0.467 (Posit Team, 2024) for each species pooling years using package *sfhotspot* v.0.8.0 (Ashby 2023), and the Getis-Ord Gi* (Gi*) statistic [64]. The hotspot method detects spatial clusters of high and low values by comparing each observation to its surrounding neighbors. We defined neighbors using Queen’s case contiguity (i.e., grid cells that share at least one corner or edge). The Gi* statistic assigns a z-score, which indicates whether the observed clustering significantly differs from a random spatial distribution, with positive z-scores representing hotspots and negative z-scores indicating cold spots. We considered the statistical significance of hotspots at α ≤ 0.05.

To examine spatial relationships among species, we imported the hotspots into ArcGIS Pro v. 3.3.0 (ESRI ArcGIS Pro©, Environmental Systems Research Institute Inc., Redland, CA), converted them to polygons, extracted the contour of significant hotspots (α = 0.05), and calculated the centroid coordinates. We measured the distance between centroids and quantified interspecific hotspot proportional overlap using:

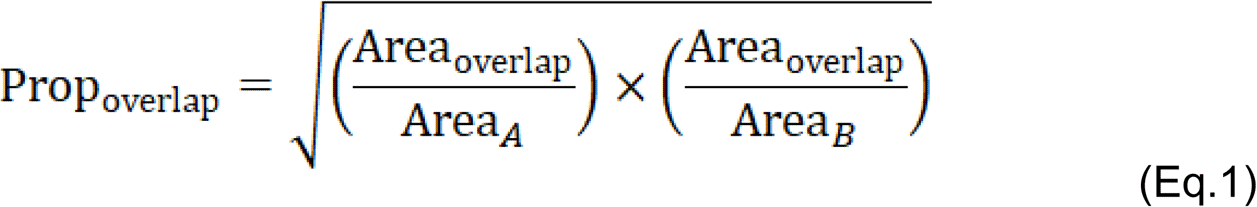

where Area_overlap_ is the area shared by the two overlapping hotspots and Area_A_ and Area_B_ are the individual hotspot areas of species A and B, respectively.

For polar bear tracks (n=3 years) and ringed seals (n=4 years), we obtained enough observations to assess temporal hotspot persistence and examine temporal changes in their spatial relationship by measuring distances between yearly centroids within each species and between species within each year. To ensure consistency of between-year comparisons, we constrained the analysis to hotspot areas within the minimum convex polygon encompassing the region common to all surveys. We quantified yearly hotspot overlap within each species using:

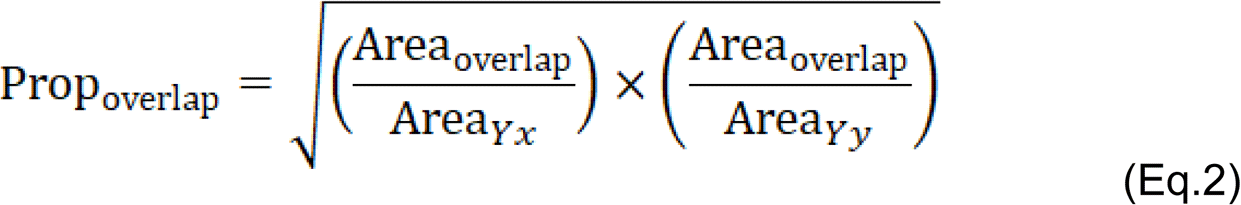

where Area_overlap_ is the area of overlap between two yearly hotspots, and Area_Yx_ and Area_Yy_ are the hotspot areas in years x and y, respectively. We assessed interannual variation in the overlap between polar bear and ringed seal hotspots by applying Equation 1 separately for each year. Descriptive statistics are provided as mean±SE.

We used directional metrics of overlap between seal-kill, hunting attempt, and species hotspots to further characterize species relationships. We quantified the proportion of polar bear and seal hotspots overlapped by seal-kill hotspot or hunting attempts, and the proportion of seal-kill or hunting-attempt hotspots overlapped by Arctic fox hotspots.

To assess relationships with sea ice, we calculated hotspot centroid distances to shore and quantified landfast vs. pack ice proportions. We obtained weekly regional ice data for Hudson Bay from the Canadian Ice Service Digital Archives (https://iceweb1.cis.ec.gc.ca/Archive/page1.xhtml; 2024). To delineate landfast ice, we selected continuous polygons with a 10/10^th^ ice concentration that were attached to the shore [65]. We calculated the maximum extent of landfast ice for each considered period (years pooled or yearly) and quantified the overlap between landfast ice and species’ hotspots using a similar equation to Eq. 1 and 2. For context, we calculated in ArcGIS Pro v. 3.3.0 the average percent of the study area covered by landfast ice, and calculated the maximum distance between landfast edge and the coast.

## Results

We surveyed 19,270 km in 498.5 hours across 39 days (Table 1) and recorded 2,412 polar bear tracks, 444 Arctic fox tracks, 697 ringed seals,148 bearded seals, 102 seal kills, and 134 hunting attempts (Table 2). Hotspots were identified for all species and seal kill locations (Fig. 2A-E). We detected no cold spots at α =0.05, thus, we focus exclusively on hotspots. Landfast ice area represented 5.0%±0.2% [4.7% - 6.0%] of the survey area weekly, and fast ice maximum width between the shore and the flaw lead was 19.2 km.

**Fig. 2.**
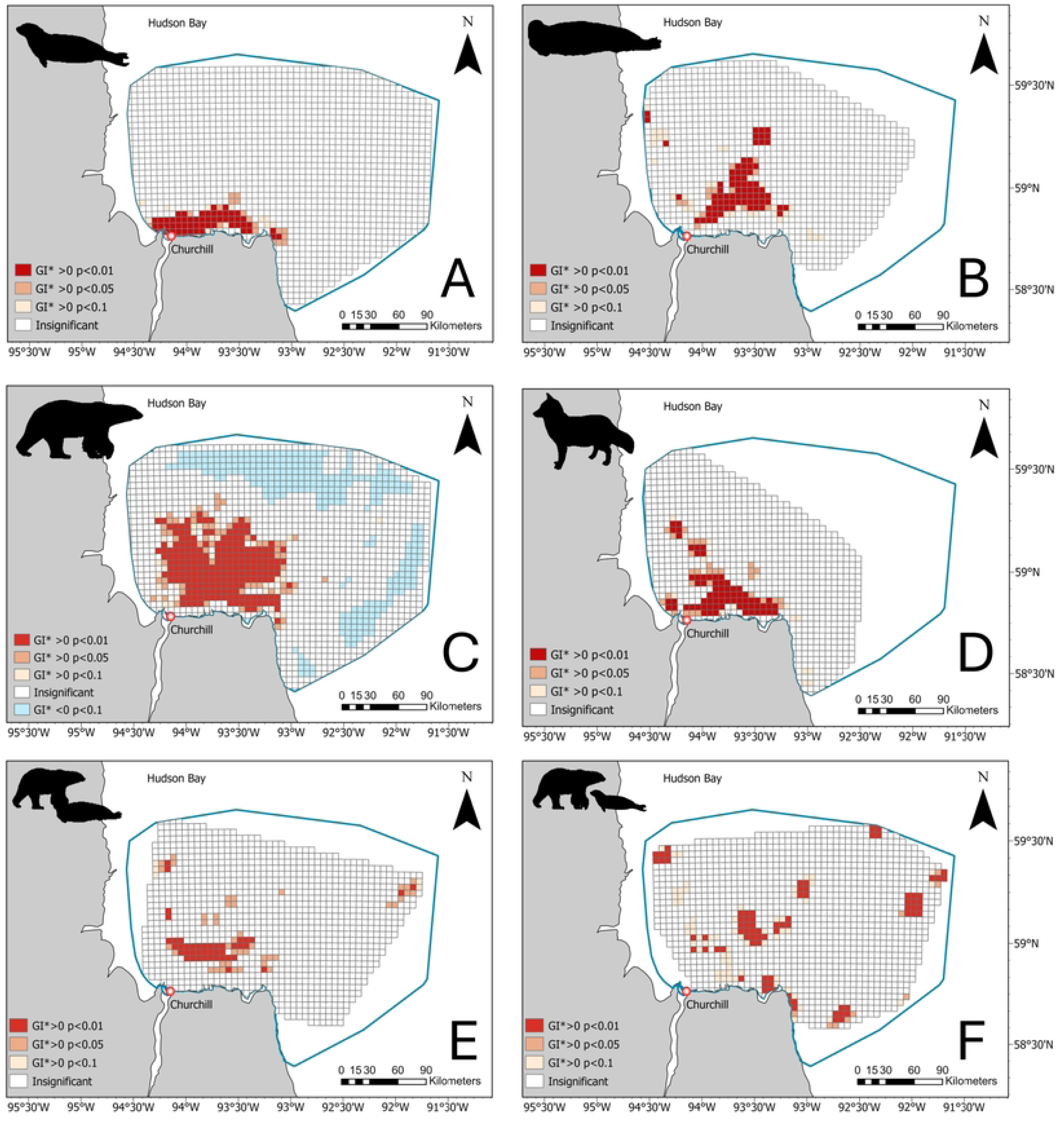
Maps of the study area in western Hudson Bay with species hotspots (A-ringed seal, B-bearded seal, C-polar bear, D-Arctic fox) and hotspot of events of seal predation by polar bears (successful: E-seal kills, or attempted F-hunting attempt) estimated with the Getis-Ord Gi* statistics. Increasing levels of red represent different levels of statistical significance. Note that we display statistically significant hotspots up to α = 0.1 but only use hotspots statistically significant at α ≤ 0.05 in subsequent spatial analyses.

**Table 2.** Raw observation counts per category (fox tracks and bear tracks were not recorded in 2019).

|  | 2019 | 2022 | 2023 | 2024 |
| --- | --- | --- | --- | --- |
| Arctic fox track |  | 192 | 207 | 45 |
| Ringed seal | 115 | 217 | 194 | 171 |
| Bearded seal | 28 | 75 | 7 | 38 |
| Polar bear hunting attempt | 17 | 42 | 36 | 39 |
| Polar bear track |  | 653 | 970 | 789 |
| Seal kill | 6 | 41 | 25 | 30 |

### Species hotspots

Pooled across all years, the ringed seal hotspot was near shore, with a centroid 3.3 km from the coast. It extended mostly west-east, covering 788 km^2^ (Fig. 2A). Yearly hotspots varied in proximity to shore, with centroid distances to shore ranging between 5.1 km in 2024 and 16.7 km in 2022 (mean = 9.9±2.9; S1 Table). Yearly-hotspot overlap proportion averaged 30% ± 5%, ranging from 14% (between 2019 and 2022) to 43% (between 2023 and 2024, when hotspots were mostly near shore; S1 Fig.). Mean interannual centroid distance was 23.4±4.9 km, ranging from 11.0 km between 2023 and 2024 to 38.2 km between 2019 and 2023 (S1 Fig.). The year-pooled hotspot was 70% landfast ice, with yearly values ranging from 26% in 2022 to 81% in 2024.

Pooled across all years, the bearded seal hotspot was mostly located offshore (Fig. 2B), with a centroid 23.3 km from shore. It covered 940 km^2^ and extended northward mostly over the pack ice, which represented 0.95 of the total hotspot area.

Pooled across all years, the polar bear hotspot extended over a large portion of the survey area, spanning 2873 km^2^, mostly on pack ice, which represented 88% of the total area (Fig. 2C) and ranged yearly between 85% (2023) and 92% (2022). The centroid was 25.2 km from shore, with yearly centroid distances to shore ranging from 21.4 to 25.4 km. Yearly hotspot overlap averaged 40% ± 6%, ranging from 28% (between 2022 and 2024) to 48% (between 2022 and 2023). Mean interannual centroid distance was 12.6±2.9 km, ranging from 11.0 km between 2023 and 2024 to 38.2 km between 2019 and 2023 (S1 Fig.).

Pooled across all years, the Arctic fox hotspot covered most of the ice near shore but extended onto the pack ice to the northwest (Fig. 2D). The centroid was 14.7 km from shore. Landfast ice comprised 38% of the total hotspot area, which was 1065 km^2^.

### Species relationships

The largest overlap between year-pooled hotspots occurred between Arctic foxes and ringed seal at 50%, while the smallest overlap occurred between ringed seals and bearded seals at 18% (Table 3). The polar bear hotspot overlapped with all three other species. The percentage overlap was 49% with Arctic foxes, 49% with bearded seals, and 30% with ringed seals. The Arctic fox and bearded seal hotspots had an intermediate overlap of 31% (Fig. 3).

**Fig. 3.**
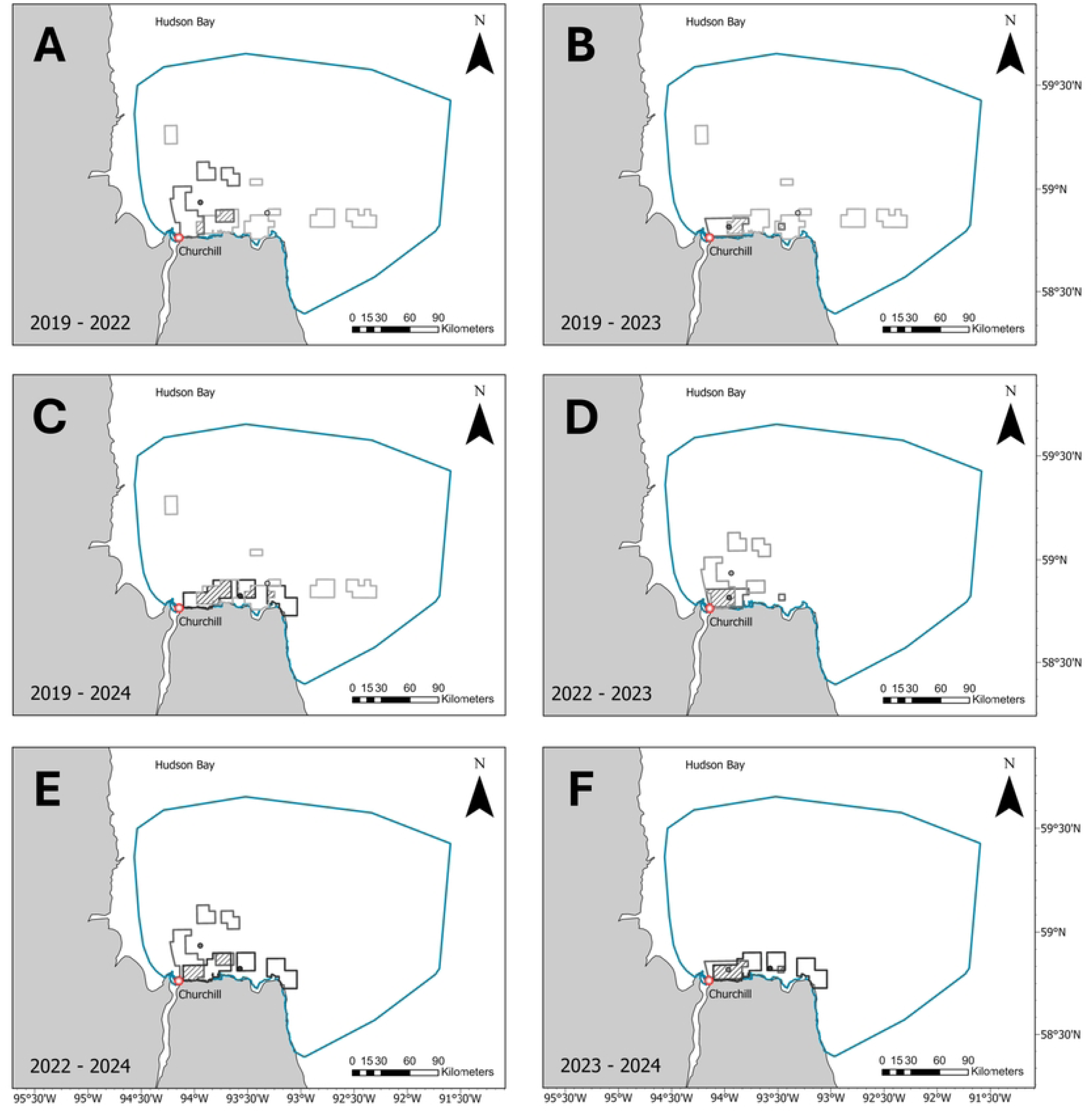
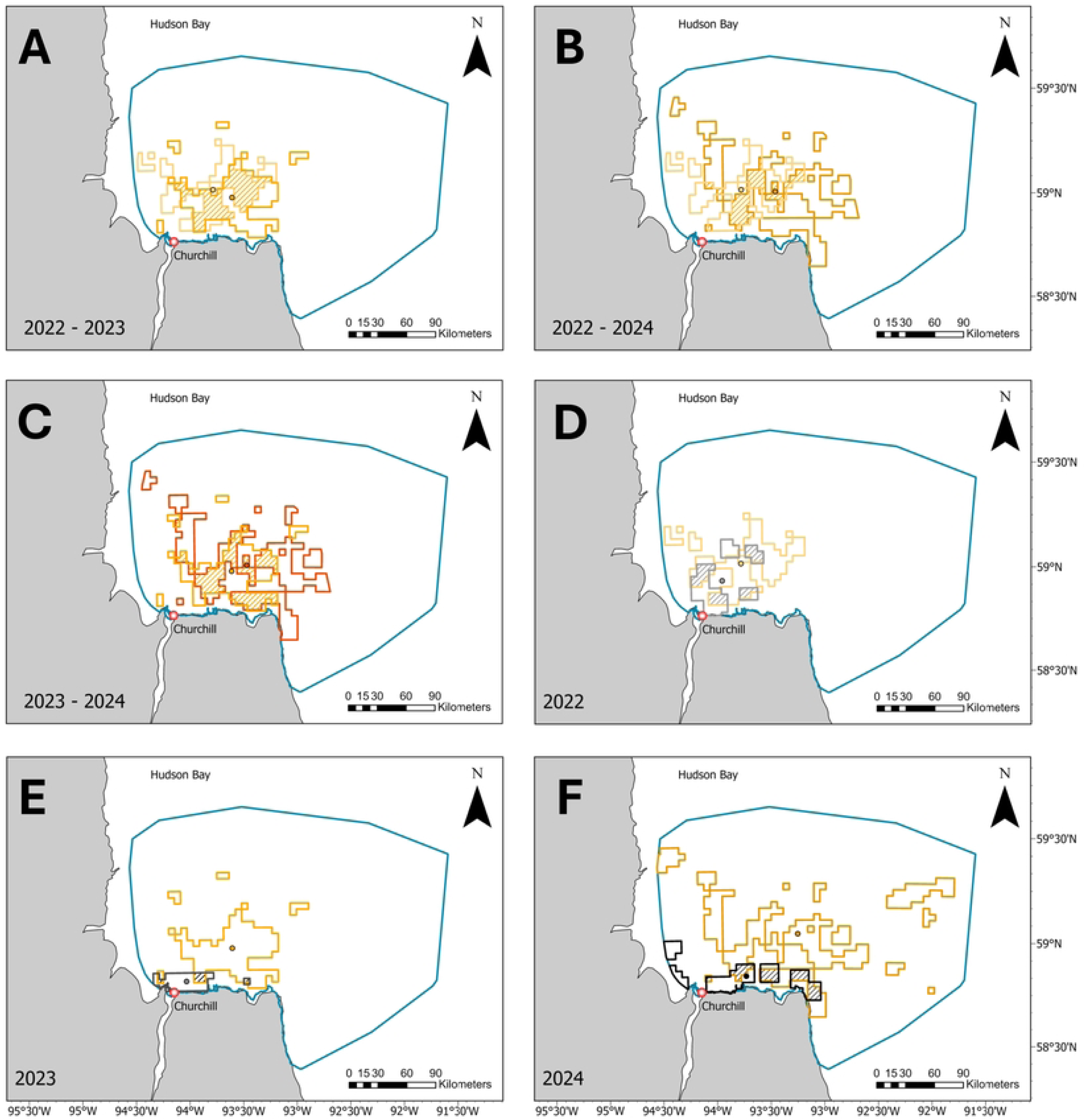
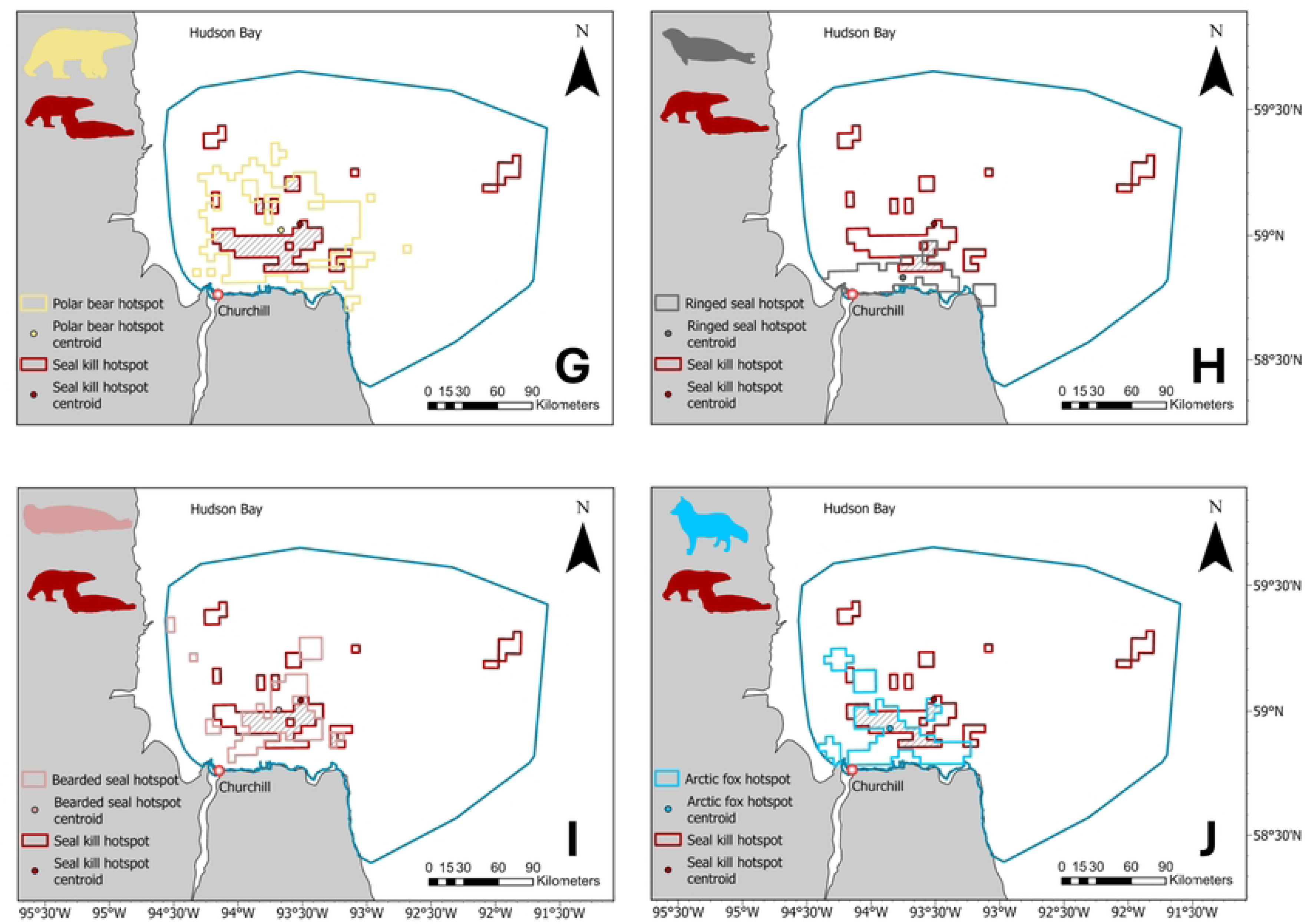
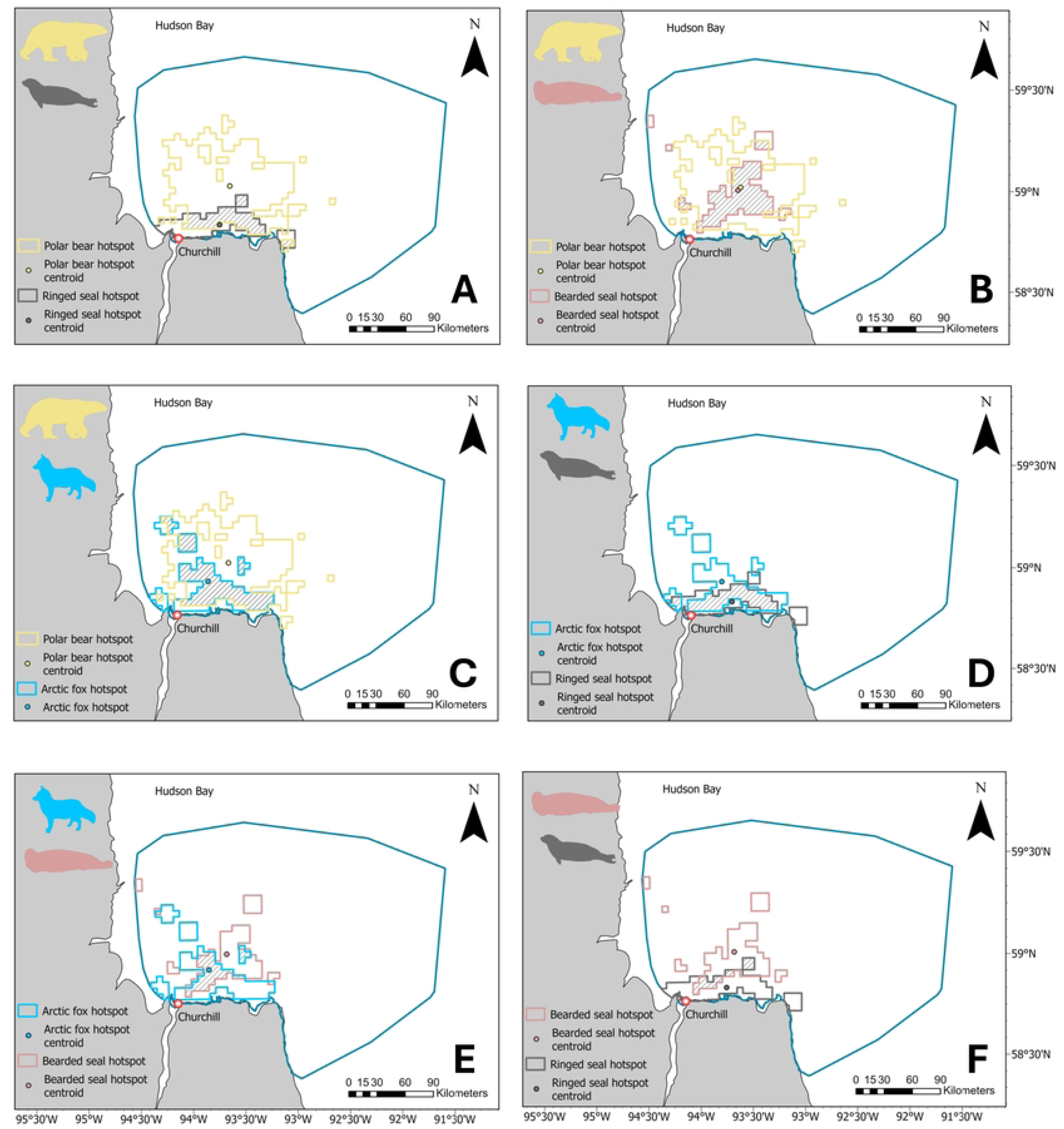
Hotspots, hotspot centroids, and hotspot overlap between species pairs (Arctic fox in blue, polar bears in yellow, ringed seals in grey, and bearded seals in pink) in the study area near Churchill, Manitoba. Hotspots were calculated using the Getis-Ord Gi* statistic. We extracted the hotspot area statistically significant at α ≤ 0.05 to produce the overlaps.

**Table 3.** Comparison of hotspot size per species, interspecific overlap between hotspot pairs, and corresponding proportion of overlap. While overlap between species is a mutual calculation of overlap, seal kill and hunting attempt overlap are directional (proportion of seal-kill or hunting attempt hotspot area overlapping polar bear and seal species hotspot area, and Arctic fox hotspot area overlapping seal-kill or hunting attempt hotspot).

| Species A | Species B | Area A (km<sup>2</sup>) | Area B (km<sup>2</sup>) | Overlap area (km<sup>2</sup>) | Overlap proportion |
| --- | --- | --- | --- | --- | --- |
| Arctic fox | Ringed seal | 1065 | 788 | 459 | 0.50 |
| Polar bear | Bearded seal | 2873 | 940 | 799 | 0.49 |
| Polar bear | Arctic fox | 2873 | 1065 | 854 | 0.49 |
| Arctic fox | Bearded seal | 1065 | 940 | 311 | 0.31 |
| Polar bear | Ringed seal | 2873 | 788 | 450 | 0.30 |
| Bearded seal | Ringed seal | 940 | 788 | 159 | 0.18 |
| Seal kill | Bearded seal | 898 | 940 | 333 | 0.35 |
| Arctic fox | Seal kill | 1065 | 898 | 316 | 0.35 |
| Seal kill | Polar bear | 898 | 2873 | 651 | 0.23 |
| Seal kill | Ringed seal | 898 | 788 | 117 | 0.15 |
| Hunting attempt | Bearded seal | 941 | 940 | 162 | 0.17 |
| Arctic fox | Hunting attempt | 1065 | 941 | 122 | 0.13 |
| Hunting attempt | Polar bear | 941 | 2873 | 372 | 0.13 |
| Hunting attempt | Ringed seal | 941 | 788 | 59 | 0.07 |

Both the seal-kill and the hunting-attempt hotspots (years pooled) were dispersed, primarily on pack ice, which represented 93% and 88%, respectively, of the total hotspot area (Fig. 2). The centroids were 29.6 km (seal kill) and 35.0 km (hunting attempt) from shore. The seal-kill hotspot overlapped 23% of the polar bear hotspot (Fig. 3G), only 15% of the ringed seal hotspot (Fig. 3H), and 35% of the bearded seal hotspot (Fig. 3I). The hunting attempt hotspot overlapped 13% of the polar bear hotspot, 7% of the bearded seal hotspot, and 17% of the ringed seal hotspot. The Arctic fox hotspot overlapped 35% of the seal-kill hotspot (Fig. 3J) and 13% of the hunting attempt hotspot.

Ringed seal and polar bear hotspot overlap varied interannually with a minimum of 9% in 2023 with a distance between centroids of 27.4 km, and a maximum of 34% in 2022 with a distance between centroids of 12.8 km. The proportional overlap in 2024 was 20% with a distance between centroids of 21.2 km (S2 Fig.).

## Discussion

Spatial overlap between predators and prey underpins their interactions, driving changes in demographic rates for both predator and prey populations, which ultimately induce cascading effects in community structure. Using direct and indirect signs of presence, we identified spatial hotspots for four ice-associated species. As we predicted, both polar bear and Arctic fox space use matched the distribution of food resources (seals or polar bear kills), while ringed seals and bearded seals showed low spatial overlap. We found that polar bears in our study had a stronger spatial association with bearded seals, compared to ringed seals. Furthermore, patterns of sea ice use by these four species were consistent with other studies [19,47,57,58].

Ringed seals mostly hauled out in nearshore areas with some interannual variation. While in 2019 and 2022 their hotspot occurred mainly on pack ice or included large areas offshore, in 2023 and 2024 they concentrated in a narrow band along the shore on the landfast ice. Furthermore, the centroid of the hunting attempt hotspot, representing predatory behavior of bears on ringed seals and therefore reflecting the distribution of ringed seal lairs, was located 35 km from the coast in pack ice, with minimal overlap of the (hauled-out) ringed seal hotspot (17%). Across their range, ringed seals typically prefer shallow waters, relatively high ice cover (40%-80%), stable and consolidated ice, and ice features that accumulate snow to build lairs [36,66–69].

They, however, display intrapopulation variability in habitat selection [36,45,70]. Notably, they can breed in the pack ice or give birth in the open when snow accumulation is too low [36,58]. In western Hudson Bay, ringed seals haul out on both landfast and pack ice, but occur in lower density on the pack ice [57]. Hudson Bay is a uniformly shallow continental shelf [62], and thus, likely offers abundant suitable ringed seal habitat, unlike the Beaufort Sea, where the bathymetry is more variable and seals are less likely to be homogeneously distributed [46]. Our findings of hauled-out ringed seals and hunting attempts at lairs reflect different seal behaviors and highlight the use of both pack ice and landfast ice in Hudson Bay, but potentially for different requirements. Our findings of ringed seal birth lairs in the pack ice indicate reproduction on this type of ice, as observed in other Arctic regions [46,58]. Habitat use could also differ by age class [45].

In contrast to ringed seals, bearded seals primarily used active pack ice, with their hotspot centroid located farther offshore, consistent with observations across the Arctic [71–73]. Consequently, the spatial overlap between hauled-out ringed seals and bearded seals hotspots was low, reflecting their different habitat preferences. Each winter, a major flaw lead forms along the coast of western Hudson Bay (including in the Churchill area), as northwestern winds push the pack ice away from the landfast ice [74]. The presence of this flaw lead enhances area suitability near the Churchill coastline for bearded seals [75].

The proportion of landfast ice in polar bear hotspots was generally low but varied considerably between years. In 2022, landfast ice accounted for 8% of the habitat within the polar bear hotspot, a percentage that nearly doubled in 2023. Given the limited variation in landfast-ice coverage within the study area, these shifts mirrored the changes in landfast ice proportion within ringed seal hotspots, consistent with their strong predator-prey relationship. However, polar bears are rarely successful when hunting adult ringed seals on landfast ice [46,63], which may explain the limited spatial overlap observed between polar bears and hauled-out ringed seals, and between the seal-kill hotspot and the ringed seal hotspot. In contrast, the bearded seal hotspot overlapped substantially with the polar bear hotspot (49%), and 35% of its area was covered by the seal-kill hotspot —highlighting the importance of bearded seals for polar bear. These findings suggest that ringed seals were not the sole focus of polar bear predation. Although bearded seals are an important secondary prey, particularly in Hudson Bay [23], the low overlap of the hunting attempt hotspot and the polar bear hotspot was unexpected, because ringed seals are the primary prey of polar bears across their range [22,23].

Polar bears select areas that maximize kill biomass rather than kill frequency alone [76]. In our study, the low spatial overlap between polar bears and hauled-out ringed seals, and high overlap between polar bear tracks and hunting activities (i.e., hunting attempts and seal kills) and bearded seals collectively suggest that bears prioritized areas with greater access to bearded seals, while still hunting ringed seals. These spatial patterns support earlier findings from the Beaufort Sea [76] and indicate that in spring, polar bears may shift their primary focus from ringed seals to bearded seals — a potentially widespread foraging strategy during the hyperphagic period. By late April to early May, although ringed seal pups remain available, their pupping peak has passed [49], while bearded seals start pupping [72], increasing overall prey availability. Under such conditions, the higher energetic cost of hunting the larger and scarcer bearded seals becomes worthwhile to maximize net energy gain, consistent with optimal foraging theory [77,78]. Selecting for pack ice may therefore represent an optimal strategy, granting access to both abundant, predictable prey with lower energetic value (i.e., ringed seal pups [79]) and scarcer but higher-yield prey (i.e., bearded seals [72,76]). Although prey abundance and catchability are often treated as competing hypotheses, some predators use hunting strategies that account for both [80,81] — a pattern that may also apply to polar bears during hyperphagia.

As predicted, the Arctic fox hotspot overlapped strongly with the polar bear (49%) and seal-kill hotspots (35%), underscoring their commensal relationship with polar bears and the importance of bear kills as a food source. Arctic foxes also showed high overlap with hauled-out ringed seals (50%), possibly reflecting predation on pups. However, since our surveys occurred after peak pupping, this association may instead reflect the role of the flaw lead as a barrier, concentrating fox activity along the landfast ice as they search for passage to the pack ice, where most bear hunting activity occurs. The low overlap with the hunting attempt hotspot suggests that foxes use the sea ice opportunistically when carcasses are available, possibly locating them from considerable distances (up to 40 km at least [82]). Anecdotally, Arctic fox tracks sometimes follow tracks of larger bears, suggesting they may maximize their chances of finding carcasses by cueing on signs from bears with higher hunting success (Derocher, pers. obs.). The Arctic fox hotspot also overlapped with the bearded seal hotspot more than expected (31%), likely due to the spatial overlap between polar bears and bearded seals. Given the size of bearded seal pups at weaning [83,84], direct predation by foxes is unlikely. Instead, foxes’ scavenging activity likely creates an indirect spatial association with bearded seals.

Arctic foxes can use sea ice extensively. For example, an Arctic fox collared in Churchill traveled nearly 5,200 km over four months on Hudson Bay’s ice (one location per day [34]). Similar long-distance movements across the Arctic [85–87] highlight their reliance on sea ice. Some foxes may even include large proportions of (landfast) sea ice in their home range during seal pupping season [34]. In our study, the Arctic fox hotspot was centered nearly 15 km offshore and comprised a higher proportion of pack ice than landfast ice. While Arctic foxes’ nearshore habitat use may reflect their vulnerability if the flaw lead opens or fidelity to their terrestrial home range, their heavy use of the pack ice highlights their adaptation to navigate this high-risk high-reward habitat.

Studying ice-associated species is challenging due to the inaccessibility of their habitat. While satellite telemetry has significantly advanced our understanding of Arctic food web dynamics, it rarely provides insights on multiple species and their spatial overlap unless each is tracked simultaneously [59,88]. Systematic collection of incidental observations of direct and indirect signs of animal presence can provide a valuable alternative for studying rare or cryptic species at low cost [46,89,90]. A limitation of the method is the inability to control detection bias, such as the greater visibility and persistence of larger carcasses [91]. Smaller ringed seal carcasses were likely consumed faster than those of bearded seals, reducing their detectability.

Consequently, more polar bear and Arctic fox tracks would be expected to travel to longer-lasting carcasses [32] potentially leading to an underestimation of ringed seal predation. While we advise caution in interpreting our findings, both spatial coverage and intensity were extensive, and the data were standardized. With careful interpretation, we believe that this cost-effective method provides a reasonable representation of true spatial patterns.

### Conservation implications

Understanding how ice-associated species use space and interact is essential for informing targeted conservation. Our cost-effective survey approach offers valuable insight into species’ spatial patterns at a time when only 8.4% of the global ocean (MPAtlas.org) and 5.2% of the Arctic marine area [92] benefit from some level of protection. For example, given the commercial significance of Churchill harbor, our findings could guide efforts to mitigate the negative effects of increasing human activity by informing the creation of marine protected areas. Species interactions, such as trophic interactions, competition, or pathogen transmission, play a central role in maintaining ecosystem balance. Thus, changes in one species’ space use or relative abundance may generate far-reaching effects for both the marine and terrestrial Arctic ecosystems. Our study highlights the importance of considering these dynamics in conservation planning to ensure effective outcomes, particularly in a rapidly changing Arctic.

## Acknowledgements

CWR is grateful for postdoctoral financial support from Polar Bears International and MITACS. Research support was provided by the Banrock Station Environmental Trust, Canadian Association of Zoos and Aquariums, Canadian Wildlife Federation, Care for the Wild International, Churchill Northern Studies Centre, Earth Rangers Foundation, Environment and Climate Change Canada, Hauser Bears, Isdell Family Foundation, Offield Family Foundation, Zuest Family Foundation, Kansas City Zoo, Manitoba Agriculture and Resource Development, Natural Sciences and Engineering Research Council of Canada, Parks Canada Agency, Pittsburgh Zoo Conservation Fund, Polar Bears International, Polar Continental Shelf Program of Natural Resources Canada, Quark Expeditions, San Diego Zoo Wildlife Alliance, Schad Foundation, Wildlife Media Inc., and World Wildlife Fund Canada. We thank Brooke Biddlecombe, Angus Derocher, Sean Headland, Natasha Klappstein, Ryan Mutz, Megan Owen, Nicholas Paroshy, Peter Thompson, Toshio Tsubota, and Yoshiko Torii for their contribution in the field.

## Supporting Information

**S1 Table.** Flight distance in km and duration in minutes per ordinal day per year (days in grey are non-survey flights and were not included in the sample effort).

**S1 Fig.** Yearly hotspots and hotspot centroids of ringed seal with interannual overlaps. In each frame, the lighter grey represents the older year. Hotspots were calculated using the Getis-Ord Gi* statistic within the area common to the 2 years compared. We extracted the hotspot area statistically significant at α ≤ 0.05 to produce the overlaps.

**S2 Fig.** Yearly hotspots and hotspot centroids of polar bears with interannual overlaps (A, B, C). In each frame, the lighter yellow represents the older year. Yearly overlap between polar-bear and ringed-seal hotspots (D, E, F). Seals are depicted in grey shades and polar bears in yellow shades. Hotspots were calculated using the Getis-Ord Gi* statistic. We extracted the hotspot area statistically significant at α ≤ 0.05 to produce the overlaps.

